# AcaFinder: genome mining for anti-CRISPR associated genes

**DOI:** 10.1101/2022.08.29.505781

**Authors:** Bowen Yang, Jinfang Zheng, Yanbin Yin

## Abstract

Anti-CRISPR (Acr) proteins are encoded by (pro)viruses to inhibit their host’s CRISPR-Cas systems. Genes encoding Acr and Aca (Acr associated) proteins often co-localize to form *acr-aca* operons. Here, we present AcaFinder as the first Aca genome mining tool. AcaFinder can: (i) predict Acas and their associated *acr-aca* operons using guilt-by-association (GBA); (ii) identify homologs of known Acas using an HMM (Hidden Markov model) database; (iii) take input genomes for potential prophages, CRISPR-Cas systems, and self-targeting spacers (STSs); and (iv) provide a standalone program (https://github.com/boweny920/AcaFinder) and a web server (http://aca.unl.edu/Aca). AcaFinder was applied to mining over 16,000 prokaryotic and 142,000 gut phage genomes. After a multi-step filtering, 36 high-confident new Aca families were identified, which is three times of the 12 known Aca families. Seven new Aca families were from major human gut bacteria (Bacteroidota, Actinobacteria, Fusobacteria) and their phages, while most known Aca families were from Proteobacteria and Firmicutes. A complex association network between Acrs and Acas was revealed by analyzing their operonic co-localizations. It appears very common in evolution that the same *aca* genes can recombine with different *acr* genes and *vice versa* to form diverse *acr-aca* operon combinations.

**Importance:** At least four bioinformatics programs have been published for genome mining of Acrs since 2020. In contrast, no bioinformatics tools are available for automated Aca discovery. As the self-transcriptional repressor of *acr-aca* operons, Aca can be viewed as anti-anti-CRISPRs, with a great potential in the improvement of CRISPR-Cas technology. Although all the 12 known Aca proteins contain a conserved Helix-Turn-Helix (HTH) domain, not all HTH-containing proteins are Acas. However, HTH-containing proteins with an adjacent Acr homologs encoded in the same genetic operon are likely Aca proteins. AcaFinder implements this guilt-by-association (GBA) idea and the idea of using HMMs of known Acas for homologs into one software package. Applying AcaFinder in screening prokaryotic and gut phage genomes reveals a complex *acr-aca* operonic co-localization network between different families of Acrs and Acas.

## Introduction

Viruses and mobile genetic elements (MGEs) have been in a constant arms race with their prokaryote hosts for billions of years (1). To prevent viral infections, prokaryotes have evolved various anti-viral defense systems, e.g., CRISPR-Cas, RM (restriction modification), TA (toxin and anti-toxin), BREX (Bacteriophage Exclusion) systems (2). To survive, viruses also evolved various anti-defense strategies (3). Among the known anti-defense systems, anti-CRISPRs have received the greatest attention due to their applications in developing more controllable and safer genome editing tools, e.g., CRISPR-Cas9 (4).

Anti-CRISPR (Acr) proteins were first discovered in 2013 from Pseudomonas phages and prophages (5). They were found to successfully protect invading phages by inhibiting host’s CRISPR-Cas systems. These published Acrs are often encoded by (pro)viral genetic operons that also contain genes of anti-CRISPR associated (Aca) proteins in the downstream of acr genes (5-7). As of now, a total of 98 Acr proteins have been experimental characterized; however, they do not share any significant sequence similarity and thus form 98 Acr protein families. Most Acrs do not have conserved Pfam domains and thus have few sequence homologs in the databases. In contrast, 12 Aca protein families have been defined and all of them contain a conserved Helix-Turn-Helix (HTH) domain, commonly found in DNA-binding proteins. Because Aca is more conserved and often co-exist with Acrs in operons, many of the 98 Acrs were actually identified by the guilt-by-association (GBA) idea followed by experimental characterization, i.e., sequence similarity search of aca (HTH-containing) genes first and then search for acr genes in the genomic operons of (pro)viruses.

Recent studies have shown that at least three Aca families negatively regulate Acr expression (8-11). These Acas bind to the promoter regions of the *acr-aca* operons via their HTH domain, leading to transcription repression of the operons (9, 10). This is reminiscent of the type II toxin and anti-toxin (TA) systems, where the anti-toxin protein is also an HTH-containing transcriptional repressor of the toxin. Therefore, Aca proteins could be viewed as anti-anti-CRISPRs, with a great potential in the calibration of CRISPR-Cas technologies by modulating the Acr modulators.

In the past three years, bioinformatics tools have been developed to aid in the discovery of novel Acrs. These include tools as automated softwares screening query genomes or ranking/scoring query proteins for Acr candidates, e.g., AcRanker, AcrFinder, PaCRISPR (12-14). There are also online databases (Anti-CRISPRdb, AcrDB, AcrHub, AcrCatalog) for experimentally verified Acrs, their homologs, and machine learning predicted Acrs (15-18). In contrast, no automated bioinformatics tools are available for Aca discovery. This might be because Acas have HTH domains and are easier to identify. However, not all HTH-containing proteins are Acas. In fact, there are 328 HTH-related Pfam families forming the HTH clan (a higher classification level than family) due to shared distant evolutionary homology. This makes the HTH clan (Pfam clan ID: CL0123) the largest and probably also one of the most conserved clan in the Pfam database. Therefore, the HTH sequence space is much larger than the Aca sequence space, and finding HTH-contain proteins does not mean that Acas are found.

Here we report the first software package, AcaFinder, to allow automated genome mining for reliable Acas. Two approaches are implemented to more confidently identify Acas given a query genome or metagenome assembled genome. The first approach is based on guilt-by-association (GBA), meaning that we identify homologs of known Acrs first and then search for HTH-containing proteins in the acr gene neighborhood. The rational is that, HTH-containing proteins are more likely to be real Acas if they are located in the same genetic operons as known Acrs or their homologs. The second approach is to build an HMM (hidden markov model) database using a training data of the 12 known Aca families, and then search for Aca homologs with this AcaHMMdb instead of Pfam HTH HMMs. In addition to the two implemented approaches, AcaFinder also calls a CRISPR-Cas search tool (CRISPRCasTyper), a prophage search tool (VIBRANT), and an in-house Self-targeting spacer (STS) searching process, providing users with detailed information vital to the assessment of Aca predictions (19-22).

Features of AcaFinder include: (i) identify both potential Acas and their associated *acr-aca* operons; (ii) identify Aca homologous proteins using the AcaHMMdb; (iii) provide potential prophage regions, CRISPR-Cas systems, and STSs in the query genome; (iv) provide a standalone software package that can run locally and a user-friendly web interface. The webserver generates graphical representations of identified *acr-aca* operons with associated CRISPR-Cas, prophage, and STS information in terms of genomic context. Lastly, using AcaFinder, we have screened for potential Aca proteins in the RefSeq prokaryote genomes and in the gut phage database (GPD) (23-26). We have performed a phylogenetic analysis to study the sequence diversity and taxonomic distribution of a group of highly confident Aca predictions.

## Materials and Methods

### Build the AcaHMMdb

For Aca homology search, we built 12 Hidden Markov Models (HMMs) corresponding to the 12 published Aca protein families with the following steps (**Fig. 1A**):

**Fig. 1.**
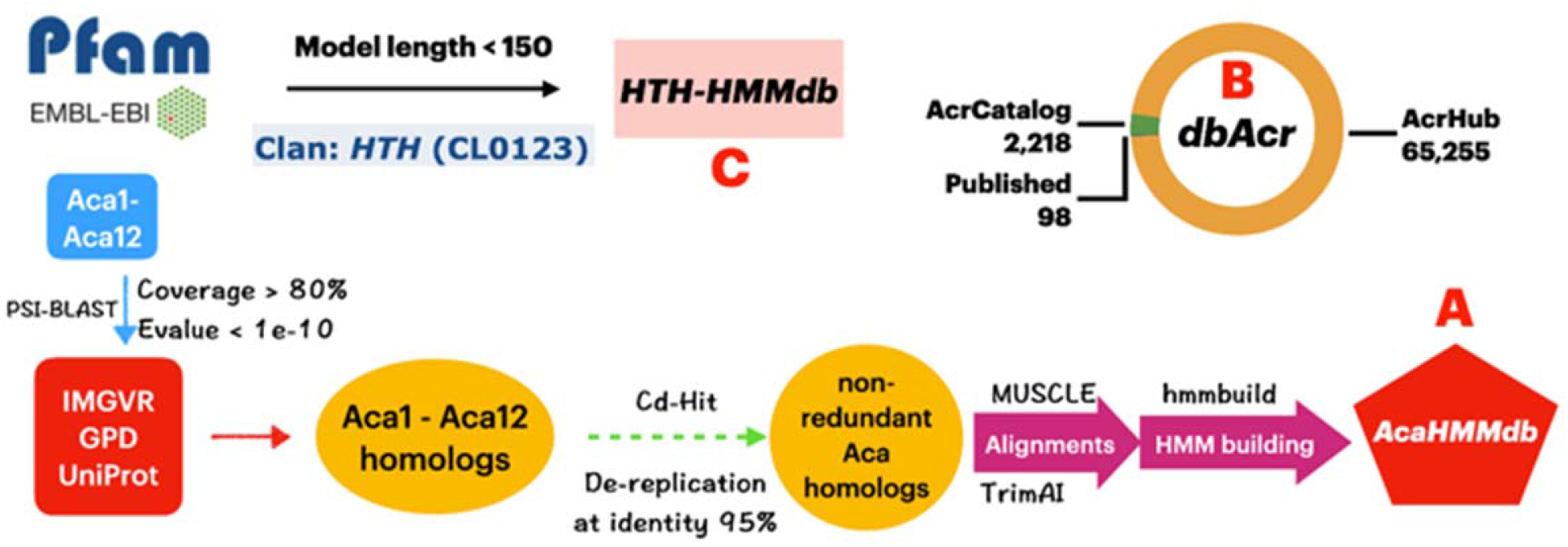
Three databases are built within AcaFinder. (A) The workflow to build the AcaHMMdb. Homologs of 12 known Acas were used in the construction of 12 HMMs. (B) The data composition of the Acr database (dbAcr). (C) The making of HTH-HMM database. Pfam database was downloaded and filtered for HTH HMMs that fit the set criteria (described in main text).

i. We downloaded the UniProt (27), IMGVR (24), and GPD protein databases;
ii. We downloaded the 12 published Aca protein (Aca1-Aca12) sequences from https://tinyurl.com/anti-CRISPR (28). Aca13 is excluded as it has no HTH domain, and not validated for Aca functions. The 12 Aca sequences were used as the query to PSI-BLAST search the three protein databases. Acr protein homologs with query coverage > 80% & Evalue < 1e-10 & protein length < 150aa were considered as Aca(1-12)-like proteins;
iii. Cd-Hit (29) was used to dereplicate the Aca(1-12)-like proteins with an identity threshold < 95%;
iv. Using MUSCLE (30), 12 multiple sequence alignments were created from the dereplicated Aca(1-12)-like proteins along with the corresponding known Acas; the alignments were then trimmed with TrimAI (31). The details of the alignment trimming are provided in **Table S1**;
v. hmmbuild (32) was used to create 12 HMMs based on the 12 trimmed alignments; the 12 HMMs were combined into one file as the AcaHMMdb.

### Collect Acr sequences to form the database of Acr (dbAcr)

For GBA search of Acas, we prepared a highly confident Acr database from two sources. The first source contains protein sequences of 98 experimental characterized Acrs downloaded from https://tinyurl.com/anti-CRISPR (28). The other source includes machine learning predicted Acr sequences that are less than 200aa in length, and share no sequence similarity with the 98 known Acrs. Those include 2,218 Acr sequences from AcrCatalog (16), and 65,255 Acr sequences from AcrHub (17). More detailed information can be found on the AcrCatalog and AcrHub websites. Therefore, the dbAcr contains 98+2,218+65,255 Acr protein sequences (**Fig. 1B**).

### Build the HTH-HMM database (HTH-HMMdb and its subset HTH-HMM_strictdb)

For GBA search of Acas, we also built an HTH HMM database. An HMMER search of the 12 known Acas against the Pfam database found that all of them have their best Pfam HMM match from the HTH clan (clan ID: CL0123) (**Table S2**). Among them, 9 Acas matched HTH HMMs with “ HTH” in their Pfam family descriptions (e.g., HTH_24, HTH_3, HTH_29), and 3 Acas matched HTH HMMs without “ HTH” in their Pfam family descriptions (e.g., KORA, DUF1870, YdaS_antitoxin).

Knowing that all known Acas match the HTH HMMs, we downloaded HMMs of the Pfam HTH clan (CL0123). Only keeping HMMs with length < 150 aa (all 12 known Acas are shorter than 150 aa), we obtained in total 328 HMMs to form the HTH-HMM database (**Fig. 1C**). Another more conservative database, the HTH-HMM_strictdb, was also built with only 89 Pfam HTH HMMs that must have “ HTH” in their family descriptions (e.g., HTH_24, HTH_3). Users have the option to choose either the HTH-HMMdb or the HTH-HMM_strictdb for searching the gene neighborhood of Acr homologs (see below).

### AcaFinder workflow

Given a nucleotide sequence fna file, AcaFinder will call gene prediction tool Prodigal (33) to generate a protein sequence faa file, and an associated gff (General Feature Format) file. In addition, AcaFinder also allows users to submit their own annotation files (fna, faa, and gff files, pink box, **Fig. 2**). After initial input, the workflow splits into two routes.

**Fig. 2.**
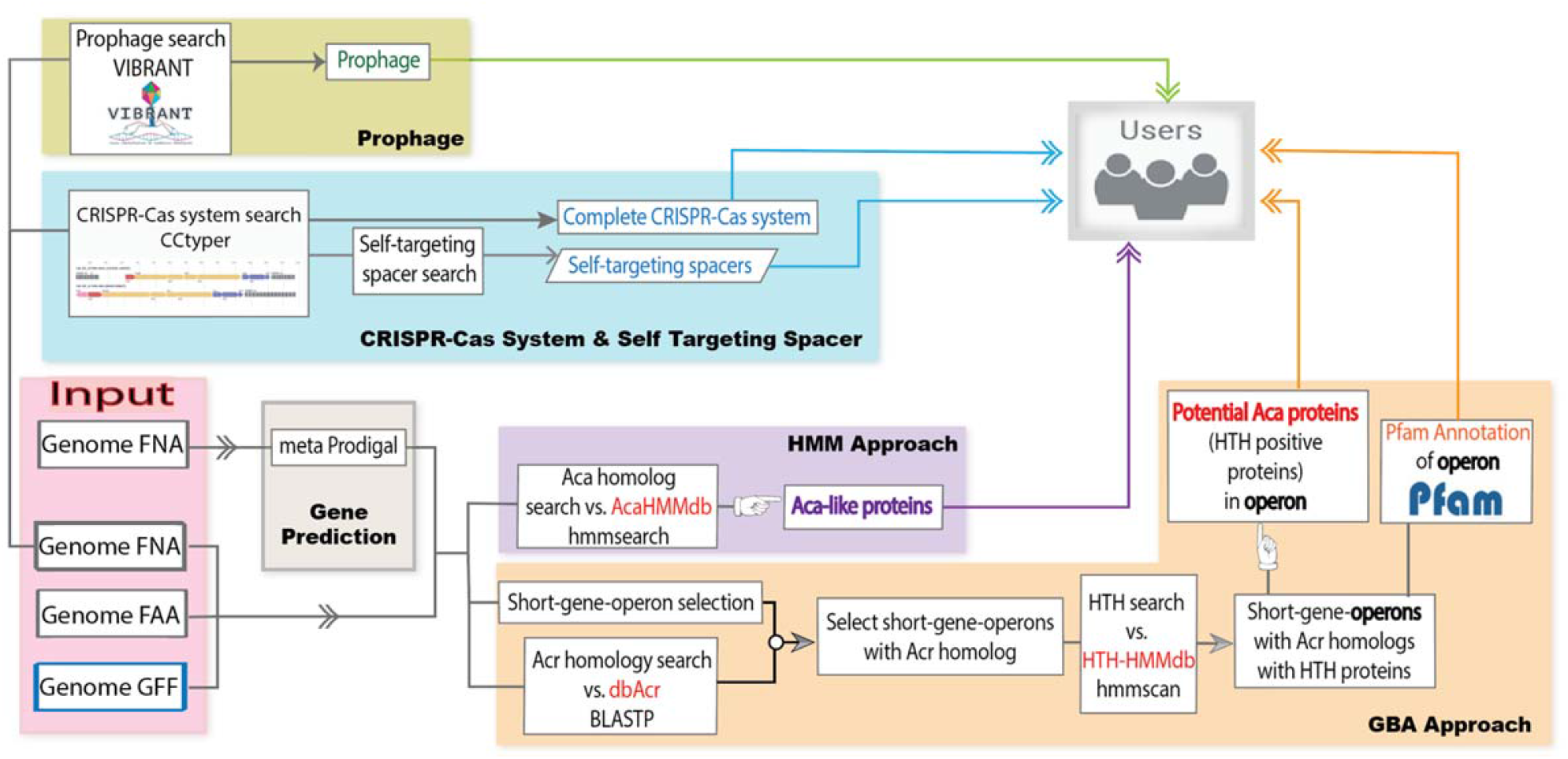
AcaFinder workflow. With provided input, AcaFinder proceeds to two Aca screening routes: (1) HMM approach to find Aca homologous proteins (purple box) and (ii) GBA (guilt-by-association) approach to find acr-aca operons (orange box), which must contain at least one Acr homolog and one HTH-containing gene (Aca candidate). Complete CRISPR-Cas along with STSs (blue box) and prophage regions (green box) are also searched from input genomic sequences (described in main text). All generated results are combined and provided to the users as tables and graphs.

One route (purple box, **Fig. 2**) uses hmmsearch with the built-in AcaHMMdb as query and user input faa file as database. Any HMM hits that passed set filters (HMM coverage > 60% & Evalue < 1e-10) will be extracted as Aca-like proteins, which will be provided as outputs to the user. The other route uses the GBA approach to find HTH-containing Aca candidates in the Acr gene neighborhood (orange box, **Fig. 2**). Compared to the simple search for HTH-containing proteins, this GBA approach considers the co-occurrence of *acr* and *aca* genes in the same genetic operons. Thus, it adds a strong constraint in the identification of highly confident novel Aca proteins. There are 3 consecutive steps:

Step 1: The input faa file will be used as query to DIAMOND blastp against the built-in dbAcr for Acr homologs (coverage > 60%, Evalue < 1e-3). Once Acr homologs are determined, the input fna file will be scanned for short-gene-operons (SGOs) by the following criteria: (i) At least one acr homologous genes in the SGO; (ii) All genes on the same strand; (iii) All intergenic distances < 250bp; and (iv) All genes have protein sequence length < 200aa (except that when the Acr homologs are homologous to known Acrs that are longer than 200aa, e.g., AcrIIIB1 [249aa], AcrVA2 [322aa]).

Step 2: Each SGO will be scanned for HTH-containing genes using hmmscan (HMM coverage

> 40%, Evalue < 1e-3) with the HTH-HMMdb or its subset HTH-HMM_strictdb as database.

Step 3: HTH-containing proteins (< 150aa) from SGOs will be output as candidate Acas. All non-Aca and non-Acr genes in SGOs will be further annotated with Pfam database using PfamScan. Information regarding each SGO, the contained Acr homolog, and candidate Acas will be provided to the user.

In addition to the identification of Acas and *acr-aca* operons, AcaFinder will also scan the input fna file for prophages, CRISPR-Cas systems, and self-targeting spacers (STSs). Bacterial genomes carrying CRISPR-Cas systems and STSs are more likely to encode Acr/Aca proteins to prevent genome self-destruction (34). Prokaryotic genome input will be scanned for complete CRISPR-Cas systems (presence of both CRISPR arrays and Cas operons) with CRISPRCasTyper (19). With the presence of complete CRISPR-Cas, blastn will be then performed with all associated CRISPR spacers as query and user’s input fna file as database (**Fig. 2**) for STSs. CRISPR-Cas and STSs information will be also provided to users independently, together with *acr-aca* operons from the previous step.

Acr/Aca genes are more likely to be found in prophages (35). In order to integrate prophage information, VIBRANT (20) will be used in the search for prophage regions within user’s prokaryotic genomic input. All discovered prophage regions will be provided to the user as a table, as well as VIBRANT’s original outputs. *acr-aca* operons or Aca-like proteins that reside in any of the identified prophages will be indicated.

AcaFinder does not report a prediction score. However, the CRISPR-Cas, STSs, and prophage output from AcaFinder will allow users to further filter the predicted Acas. We recommend high-quality Acas fit the following categories: (i) predicted by AcaHMMdb, i.e., high sequence homology to the 12 known Acas; (ii) located within a predicted prophage region; (iii) from genomes that contain complete CRISPR-Cas systems and STSs.

## Results

### Performance evaluation of AcaFinder

AcaFinder is the first computational tool for automated Aca identification. The AcaHMM homology approach (**Fig. 2**) can identify Aca-like proteins sharing significant homology to the 12 known Acas. The GBA approach is more sensitive as it will identify all short HTH-containing proteins that reside in the same genetic operons with homologs of published Acrs. To evaluate the performance of AcaFinder, we have run it on the source genomes of the 12 known Acas. For each of the 12 known Acas, the genomic sequences (fna file) as well as protein annotations (faa, gff files) were downloaded from NCBI and used as input to run the AcaFinder.

Using the GBA approach (yellow box, **Fig. 2**), all 12 known Acas can be identified (**Table 1**), giving the approach a recall of 100%. With the HMM approach (purple box, **Fig. 2**), 10/12 (83.3%) known Acas were identified using the default parameter setting (HMM coverage > 60% & Evalue < 1e-10), which is designed to reduce false positives. As expected, when more relaxed thresholds were used (coverage > 30% & Evalue < 1e-3), the two missed known Acas (Aca6 and Aca9) were indeed found back (**Table S3**).

**Table 1:** Test AcaFinder on genomes encoding the 12 known Acas.

| Genome ID | Known Aca | Length (aa) | GenBank ID | Found by AcaFinder | Total Aca candidates found* | Found in leave one out** |
| --- | --- | --- | --- | --- | --- | --- |
| GCF_000904095.1 | Aca1 | 79 | YP_007392343 | Yes | 1+4 | N+N |
| GCF_000381965.1 | Aca2 | 125 | WP_019933869.1 | Yes | 1+4 | N+N |
| GCF_001066195.1 | Aca3 | 70 | WP_049360086.1 | Yes | 3+2 | N+Y |
| GCA_002194095.1 | Aca4 | 67 | OWI97558.1 | Yes | 8+5 | N+N |
| GCF_000754765.1 | Aca5 | 60 | WP_039494319.1 | Yes | 4+2 | N+N |
| GCF_000518385.1 | Aca6 | 65 | WP_035450933.1 | Yes | 2+5 | N+N |
| GCF_001662305.1 | Aca7 | 68 | WP_064702654.1 | Yes | 3+1 | N+Y |
| GCF_001725895.1 | Aca8 | 55 | YP_009272953.1 | Yes | 2+3 | N+Y |
| GCF_004135975.1 | Aca9 | 69 | WP_129352084.1 | Yes | 2+1 | N+Y |
| GCA_004745455.1 | Aca10 | 65 | TGC30851.1 | Yes | 3+4 | Y+Y |
| GCF_000314775.2 | Aca11 | 63 | WP_009730540.1 | Yes | 12+7 | N+Y |
| GCF_000526075.1 | Aca12 | 140 | WP_231458942.1 | Yes | 18+16 | Y+Y |

\* two numbers: 1st is the Aca count from AcaHMM search, and 2nd is from GBA search. Details are provided in Table S3; \*\*two values: 1s is to indicate if the known Aca was identified from AcaHMM search (Y=Yes, N=No), and 2nd is from GBA search.

In addition to recall (the true positives divided by all positives [12]), precision is also often reported in bioinformatics tool evaluation. Precision is calculated as the true positives divided by all predictions (true positives + false positives). However, the 12 known Acas only represent a very tiny fraction of all the possible Acas that exist (36). Thus, it is not rational and acceptable to treat all other predictions as false positives, i.e., those candidate Acas from AcaHMMdb search and *acr-aca* operons from GBA search in the same genomes as the known Acas. In fact, it is observed that one genome can encode multiple types of Acrs (37-39) and Acas (16, 40, 41). In AcaFinder search results (**Table 1** and **Table S3**), all the source genomes encoding the 12 known Acas had additional Aca candidates identified. Currently, we do not know whether they are real Acas or false positives, except that they have significant sequence similarity with known Acas (E-value < 1e-10 & coverage > 60%) or are present in short gene operons (SGOs) that also encode Acr homologs. However, in most cases, the GBA gives a smaller number of Aca candidates, which may be more confident predictions given the strict SGO constraint (**Table 1**).

Although there are no other similar tools that can be compared, we sought to evaluate the performance of AcaFinder’s GBA approach using leave-one-out (LOO) experiments of the 12 known Acas. The idea of LOO is that we will remove one Aca in each experiment, more specifically, by removing all published Acrs that are known to co-localize with that Aca from the dbAcr. The co-localizations of Acr and Aca (**Fig. 3** and **Table S4**) were manually curated from literature. For example, Aca2 has been shown to co-localize with AcrIF6, AcrIF8, AcrIF9, AcrIF10, AcrIIC1, AcrIIC2, AcrIIC4, and AcrIIC5 in the literature (42-45). However, the known co-localizations are obviously incomplete, and numerous unknown *acr-aca* co-localizations remain to be characterized. In the LOO experiment of Aca2, we removed all known Acr associates and then ran through the GBA route of the AcaFinder to see if Aca2 can be identified in the source genome of Acas.

**Fig. 3.**
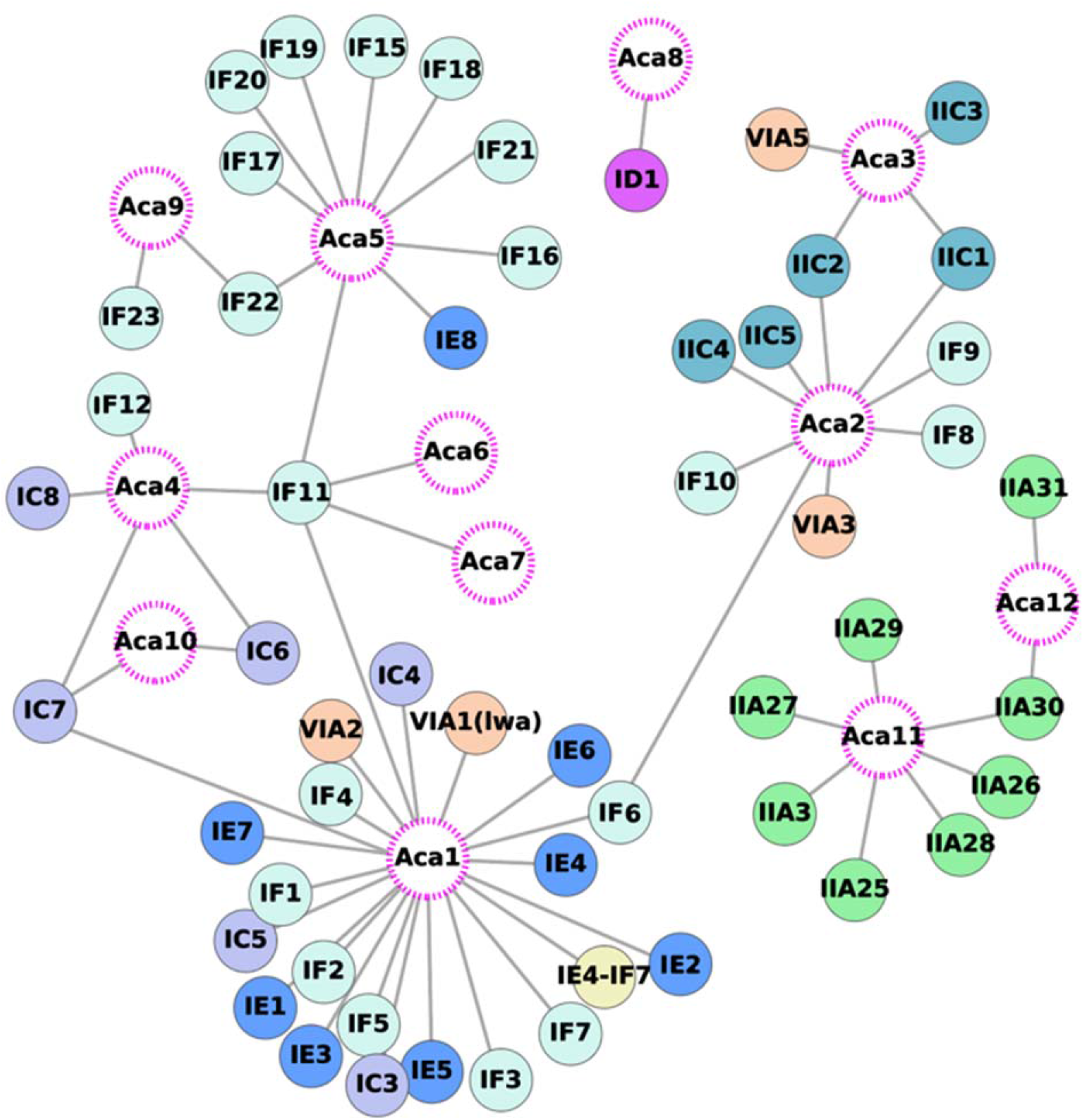
Co-localizations of known Acrs and Acas curated from literature. The 12 published Aca families are shown in open circles with pink dotted edges; Acr families are shown in filled circles colored based on type. The detailed data can be found in **Table S4**.

The result (**Table 1**) shows that 7/12 known Acas were found in the LOO experiments (recall=58.3%), while Aca1, 2, 4, 5, 6 could not to be found after removing Acrs known to co-localize with them. Note that the GBA approach has the assumption that aca genes of one family can be recruited to the gene neighborhood of different acr gene families, and *vice versa*. Therefore, if we remove all Acrs known to co-localize with an Aca, we will not find the Aca back, unless some other Acrs in dbAcr also co-localize with the Aca but such co-localizations have not been reported yet (i.e., not in **Fig. 3** and **Table S4**). Therefore, the reason that Acas1, 2, 4, 5, 6 failed to be found is because after removing Acrs known to co-localize with these Acas, no other Acrs in dbAcr can find them back. However, new Acrs are constantly being characterized. It is likely that continuously adding new Acrs in dbAcr will reveal more *acr-aca* co-localizations and improve the recall of AcaFinder in LOO experiments.

We have also run the LOO experiments to evaluate AcaFinder’s AcaHMM approach as the baseline. For each Aca, we removed its HMM from the AcaHMMdb and ran AcaFinder to see if the Aca can be found by the remaining 11 Aca HMMs. As expected, only 2 Acas were found in the LOO experiments (**Table 1**): Aca10 was identified by Aca1 HMM, and Aca12 by Aca11 HMM. Obviously, the AcaHMM approach is not good for finding novel Aca families, as between different Aca families there are very low sequence similarities.

### Utilities of AcaFinder webserver and standalone program

AcaFinder is provided as a standalone program and a webserver. Genomic sequences in fna, faa, and gff files are accepted as input. Only providing a fna file is also allowed and recommended, in which case, gff and faa files will be generated by running Prodigal (33). Genomic data of archaea, bacteria, and viruses are all allowed. By providing the Virus flag (--Virus), viral data will not run CCtyper for CRISPR-Cas scanning (STSs search also will not run), nor VIBRANT for prophage prediction.

The AcaFinder website is powered by SQLite + Django + JavaScript + Apache + HTML. A help page is available to provide users with step-by-step instructions on the usage of the webserver, along with the interpretation of the outputs. Users can submit FASTA sequences of their genomes (**Fig. S1A**). A prokaryotic genome on average takes 10 min of runtime, whereas a virus genome/contig takes 1 min. Users can skip prophage and CRISPR-Cas searches, and the job predicting Acas can finish in 1-2 min on average. The result page has tables showing information on predicted Aca proteins and associated genes within the same *acr-aca* operons of the GBA approach, and the Aca-like proteins from the AcaHMM approach (**Fig. S1B**). Prophage and complete CRISPR-Cas system information will be provided if identified, also their relationship with the predicted Acas and *acr-aca* operons will be indicated (**Fig. S1C**). Graphical representations will be generated if the input is prokaryotic genome, showing where the aca genes/operons are located on the input genome/contig(s), and how far they are from prophage regions and CRISPR-Cas systems. If any STSs were found, a link will connect the CRISPR spacer and the target region on the genome/contig(s) (**Fig. S1D**).

### Genome mining for new Aca families

A total of 15,201 complete bacterial genome, 961 archaea genomes of the RefSeq database (46) and 142,809 viral contigs of the human gut phage database (GPD) (47) were scanned with AcaFinder using the GBA approach. We used the “ --HTH-HMM_strictdb” flag for potential Aca proteins/operons. To further increase the confidence level of predictions, we limited the predicted Acas from RefSeq to genomes with complete CRISPR-Cas systems and within prophage regions. To cluster predicted Acas into potential families, Cd-Hit (29) was used with a 40% sequence identity threshold, on the basis of proteins above this threshold are more likely to share structure and function similarities (14). Cd-Hit clusters/families with a size >= 3 were selected to filter out singletons and smaller size Aca families.

After all these processes a total of 1,422 Aca families were found (**Table S5** and **Fig. S2**). We further selected Aca families that have at least one Aca member located next to homologs of the 98 experimentally characterized Acr proteins. Only 36 Aca families remained but were considered to have a very high probability being true Aca families. Combining the representative sequences of the 36 families and the 12 published Acas, a phylogenetic tree was constructed using the multiple sequence alignment the 48 protein sequences (48, 49) (**Fig. 4A**). These proteins form two major clades, and the 12 published Acas are spread across different clades of the constructed tree (**Fig. 4A**, and **Fig. S2** for a larger tree with all 1,422 Aca families). This indicates that the 12 published Aca proteins are of rather distant families, and also strengthens the point that a large sequence diversity of Acas in nature awaits to be discovered. Some of the new Aca families have large size (e.g., over 200 members, **Fig. 4B**), and the average family size is 49. Among the 48 families, 8 contain members from both RefSeq prokaryotes and GPD phages, 27 are exclusively from GPD phages, and 13 exclusively from RefSeq prokaryotes.

**Fig. 4.**
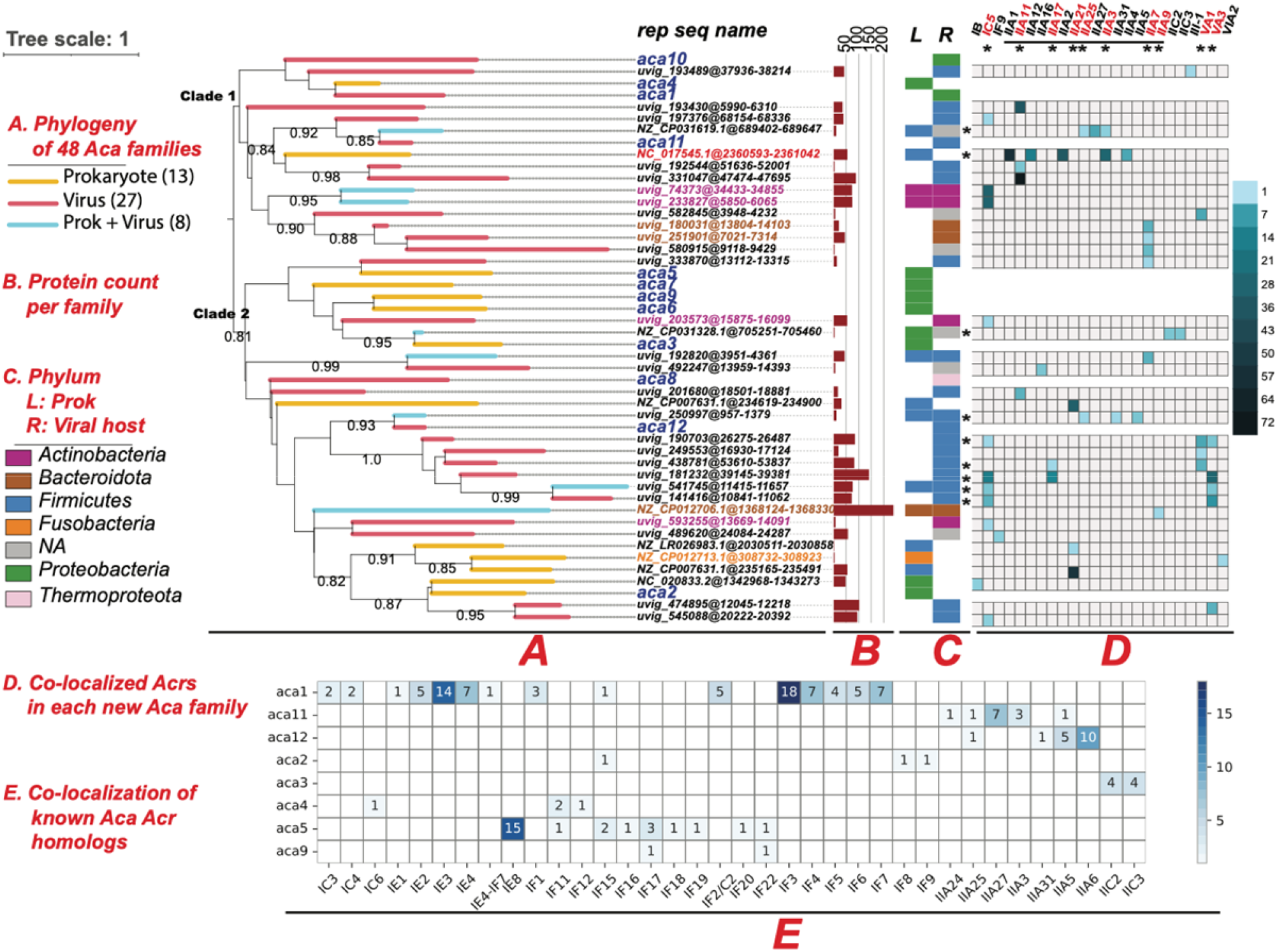
Phylogenetic and taxonomic distribution of 48 Aca families. The 36 new Aca family representatives plus 12 published Aca protein sequences were aligned with MAFFT. The phylogeny was built using FastTree and then visualized in the iTOL web server (50). (A) Branch colors represent Prokaryotic (yellow), Viral (red), or Prokaryotic + Viral (blue) protein content of each Aca family. Bootstrap values > 0.8 are shown beside the nodes; seq ID of each node are located to the right of the tree, with 12 published Aca highlighted in blue, and the predicted Acas printed in “ Contig” @” genomic location” format (e.g., NZ_LR026983.1@2030511-2030858). (B) Protein counts of each family are presented as barplot (dark red); (C) Phyla of Aca proteins are displayed in two stacked bars, with one representing prokaryotic proteins (Left), the other viral proteins (Right). For Viral proteins, the hosts’ phyla are displayed. The legend of the stacked barplot resides at the left most of the fig.; (D) The heatmap displays the member counts of the 36 Aca families that are co-localized with homologs of the 89 known Acr families (columns) in genomic operons. Rows and columns are labeled with ‘*’ if they have multiple filled cells, meaning that Aca family is co-localized with more than one Acr families, and vice versa. (E) The heatmap shows the co-localization of protein homolog counts of known Aca families and homologs of the 89 known Acr families in genomic operons of RefSeq and GPD genomes.

We have examined the taxonomic origin of the member proteins of each Aca family at the phylum level. The phylum of member proteins of RefSeq prokaryotes was plotted (L or left of **Fig. 4C**). However, most viral contigs of GPD were not assigned to a viral taxonomy group. Hence, we plotted the phylum of their hosts (R or right stripe of **Fig. 4C**), which were determined in the original paper of GPD (47). The 12 known Acas are from two phyla: Proteobacteria and Firmicutes (except Aca8 from viruses of Thermoproteota archaea), while the 36 new Aca families are distributed in three additional bacterial phyla and their phages (Bacteroidota, Actinobacteria, Fusobacteria) known to be important human gut bacteria. Note that the 4 Aca families of Actinobacteria are all co-localized with AcrIC5, and the 3 Aca families of Bacteroidota origin are all co-localized with AcrIIA7 and IIA9 (**Fig. 4D**).

We have studied the co-localizations of Acas and Acrs in the same genetic operons. First, **Fig. 3** depicts all the known *acr-aca* co-localizations extracted from literature (**Table S4**), showing that 9 (75%) of 12 known Aca families are co-localized with multiple known Acr families. Aca1, Aca2, Aca5, Aca11 have 22, 9, 10, and 7 co-localized Acr families, respectively. Some Acas are even co-localized with different Acr types (e.g., Aca1 with Acr type IE, IF, IC, and VIA). Similarly, 5 Acr families are co-localized with multiple Acas. For example, AcrIC7 is co-localized with Aca1, Aca4, and Aca10, and AcrIF11 is with Aca1 and Acas 4-7 (**Fig. 3**). Although there are many Acrs co-localizing with just one Aca, only three Acas co-localize with exclusively one Acr (Aca6-AcrIF11, Aca7-AcrIF11, Aca8-AcrID1).

In the attempt to expand these observations, a homology search against the RefSeq and GPD genomes with known Aca and Acr proteins as query was performed using blastp (coverage > 80%, Evalue < 1e-10). Only Acr and Aca homologs present in the same SGOs were considered as co-localizations, and their instances were recorded (**Fig. 4E, Table S6**). Most of the co-localizations are already known (**Fig. 3**) but there are a few new ones (e.g., Aca1-AcrIF15, Aca2-AcrIF15, Aca9-AcrIF17, Aca11-AcrIIA24, Aca12-AcrIIA5, Aca12-AcrIIA6, **Fig. 4E**). Data in **Fig. 3** and **Fig. 4E** suggest that it is very common in evolution that the same aca genes recombine with different acr genes and *vice versa* to form diverse *acr-aca* operon combinations.

We further examined the co-localizations of the 36 new Acas and known Acr homologs. We found that 9 (25%) of the 36 new Aca families are co-localized with multiple Acr families, which is different from known *acr-aca* co-localizations (**Fig. 3**). This is probably due to our stringent filtering process for the 36 new Aca families. The new Aca family “ NC_017545.1@2360593-2361042” of Firmicutes has the most types of co-localized Acrs (AcrIIA1+AcrIIA12+AcrIIA2+AcrIIA3+AcrIIA4, **Fig. 4D**). This suggests this Aca family has evolved to regulate various subtype IIA Acr subfamilies in Firmicutes. We also found that 10 Acrs are co-localized with multiple new Aca families, compared to only 5 Acrs co-localizing with multiple known Acas (**Fig. 3**). AcrIC5 is known to co-localize with Aca1 (**Fig. 3**) but co-localize with 10 new Aca families from distant taxonomic groups (**Fig. 4D**). Similarly, AcrIIA11 co-localize with 4, AcrIIA7 with 5, AcrVA1 with 4, AcrVA3 with 4 new Aca families. These diverse Aca family co-localizations indicate these Acrs flexibility in accepting different Aca families as transcriptional regulators.

## Discussion

Anti-CRISPRs (Acrs) have been under extensive studies since the discovery in 2013. These studies include various biotechnical and biomedical applications of Acrs, e.g., in genome editing, with very promising success (4, 51). Compared to Acrs, Acas are under-researched, although as anti-anti-CRISPRs, Acas also have great potentials in the development of genome editing biotechnology. In addition, due to Aca’s association with Acrs, the study of Acas can assist the continuous discovery of novel Acrs (5, 8).

AcaFinder is the first automated Aca prediction tool. A common practice in the discovery of the 12 published Acas is to search the Acr gene neighborhood for Pfam HTH-contain proteins (i.e., the GBA approach). AcaFinder implements this GBA approach and an Aca HMM approach (**Fig. 2**) as an automated computer program and webserver, and thus will assist anti-CRISPR researchers to perform more rapid genome screening for Acas.

Because there are no other similar tools that we can compare AcaFinder with, to evaluate its performance, we have conducted LOO (leave one out) experiments using the 12 known Acas. The recall of the GBA approach in LOO experiments was 58.3%. This low recall is not surprising because of the way LOO worked: removing all Acrs associated/co-localized with an Aca from dbAcr and using other Acrs to find the Aca back. GBA requires the search for Acr homologs as the first step. If the remaining Acrs in dbAcr do not have homologs in the gene neighborhood of the tested Aca, then the Aca will not be found. Therefore, the GBA approach relies on the assumption that the same Acr family can recombine with different Aca families to form *acr-aca* operons in different genomes, and *vice versa*. This assumption is supported by the observations made in **Fig. 3, Fig. 4D**, and **Fig. 4E**, which only represent a small fraction of possible *acr-aca* associations in nature. Future characterization of more Acrs, Acas, and their co-localizations to form genetic operons will undoubtedly reveal a much higher diversity of Acr and Aca combinations and help improve the power of the GBA approach in AcaFinder.

AcaFinder has the following limitations: 1) The GBA approach relies on Acr references to locate short-gene operons, thus certain Acas that do not have Acrs in proximity will be missed, and that Acas predicted may be biased towards the Acrs used for reference; 2) The Aca HMM approach relies on HMMs built from sequence alignments of known Aca homologs, thus novel Acas that share little to none sequence similarity to any known Acas will likely be missed. The drawbacks mentioned are due to the workflow design, which was intended to minimize false positives. With the continuous characterization of novel Acrs and Acas, we plan to update AcaFinder once a year to update the AcaHMMdb and dbAcr.

## Acknowledgements

We thank the lab members for helpful discussions. This work was partially completed utilizing the Holland Computing Center of the University of Nebraska–Lincoln, which receives support from the Nebraska Research Initiative.

## Funding

This work has been supported by the National Institutes of Health (NIH) awards [R21AI171952] and [R01GM140370], the United States Department of Agriculture (USDA) award (58-8042-7-089), and the Nebraska Tobacco Settlement Biomedical Research Enhancement Funds as part of a start-up grant of UNL [2019-YIN] to Y.Y.

## Supplemental materials

Table S1: Homolog sequence alignments trimming for HMM building of 12 known Acas

Table S2: PfamScan top hit of 12 known Acas

Table S3: AcaFinder performance evaluation on the genomes of 12 known Acas

Table S4: Aca associated Acrs from literature

Table S5: 1422 predicted high quality Aca families (representatives)

Table S6: Homologs of known Acrs and Acas co-localizations

Figure S1: Screenshots of AcaFinder job submission page and result page

Figure S2: Phylogeny of 1422 predicted Aca representative sequences + 12 known Aca sequences

